## Supplementary Methods, Results, Figures for "Abundant pleiotropy across neuroimaging modalities identified through a multivariate genome-wide association study"

1. *Sample description TOP, BUPGEN, MoBa……………………………………………………….2*

**Supplementary Figures**

1. *Univariate heritability estimates……………………………………………………………….7*
2. *Phenotypic and genetic correlation matrix……………………………………………………..8*
3. *Sensitivity analyses……………………………………………………………………………….9*
4. *Multivariate Manhattan plots…………………………………………………………………….10*
5. *Replication of overlapping loci and genes in ABCD…………………………………………….11*
6. *Heatmap of all univariate p-values underlying multivariate lead SNPs…………………..12*
7. *Highest univariate p-values underlying multivariate lead SNPs…………………….…………13*
8. *Single-modality and cross-modality temporal gene-expression……………………….……14*
9. *Tissue specificity of original and condFDR genes...……………………………………………15*
10. *PCA of diffusion MRI phenotypes……………………………………...……………………….16*

**Supplementary References**

### Supplementary Methods

#### Samples

We obtained individual level genotype data from three independent cohorts to predict disease liability by polygenic scores as described in the Methods section. The sample sizes and characteristics of healthy controls and schizophrenia, major depressive disorder, bipolar disorder (TOP), autism spectrum disorder (BUPGEN), and attention-deficit hyperactivity disorder (MoBa) patients are described in Supplementary Data 12.

*TOP*

The Thematically Organized Psychosis (TOP) study received ethical approval from Norwegian REC (ref. 2009/2485), Data Inspectorate (ref. 03/02051), and The Norwegian Directorate of Health (ref. 05/5821). Patients were recruited from the mental health clinics of the major hospitals in Oslo, Norway. All patients meeting diagnostic criteria were between 18 and 65 years of age and provided consent of participation. Exclusion criteria were a history of severe somatic disease interfering with brain functioning including neurological disease, history of moderate or severe head trauma, IQ < 70, and being in the initial phase of antipsychotic drug trials during recruitment. Demographic data, information about ancestry and information about current and past five years drug treatment were collected by interview and from medical records. Healthy controls were randomly selected from the national population register and included when aged between 13 and 72 years old, and in the absence of a current or previous psychiatric disorder as identified by the Primary Care Evaluation of Mental Disorders (Prime-MD) delivered by a trained research assistant[1]. Further exclusion criteria were substance use disorder, physical health condition, previous traumatic brain injury, neurological disorders, autism spectrum disorder, personal or family (1st degree relative) history of severe psychiatric disorder.

*MoBa*

The Norwegian Mother, Father and Child Cohort Study (MoBa) is a population-based pregnancy cohort study conducted by the Norwegian Institute of Public Health[2]. Participants were recruited from across Norway from 1999-2008. The women consented to participation in 41% of the pregnancies. The cohort includes approximately 114,500 children, 95,200 mothers and 75,200 fathers. The current study is based on version 12 of the quality-assured data files released for research in January 2019. The establishment of MoBa and initial data collection was based on a license from the Norwegian Data Protection Agency and approval from The Regional Committees for Medical and Health Research Ethics. The MoBa cohort is currently regulated by the Norwegian Health Registry Act. The current study was approved by The Regional Committees for Medical and Health Research Ethics (2016/1226/REK sør-øst C). All MoBa diagnoses used in the study were asserted with the corresponding diagnosis in ICD revisions 8, 9 and 10 from The Norwegian Patient Registry (NPR) registered in specialist health care services from

2008 through 2018 (using the *R* package Phenotools for extraction of the diagnostic data https://github.com/psychgen/phenotools). All subjects not diagnosed with the phenotype of interest were used as controls.

*BUPGEN*

BUPGEN is an ongoing multisite study on neurodevelopmental disorders funded by Research Council of Norway. Since 2010 more than 1500 individuals from all health regions of Norway were recruited to the study including patients under investigation or suspicion of an autism spectrum diagnosis and/or individuals carrying a genetic variant associated with autism spectrum disorder and healthy controls, the BUPGEN study thereby comprising the whole autism spectrum. In the current study only unrelated participants with a clinical diagnose of ASD according to ICD-10 (F84x) and of European decent, are included. ASD diagnoses were assigned by Norwegian specialist health services, using Autism Diagnostic Observation Schedule (ADOS) or the Autism Diagnostic Interview-Revised (ADI-R), or both as part of standard clinical evaluation. The BUPGEN study received ethical approvals from REC South East (#240277; #16858; #26141). The participants themselves or their guardians have signed a written informed consent form.

**Genotype data**

Blood and saliva samples from TOP and BUPGEN participants were used to extract DNA and genotyping was performed on Human Omni Express-24 v.1.1 (Illumina Inc., San Diego, CA, USA) at deCODE Genetics (Reykjavik, Iceland). Pre-imputation quality control was performed using PLINK1.9[3]. Variants were filtered on genotyping rate < 95%, HWE $p < 1\times10^{-4}$, a high rate of Mendel errors in eventual trios or significant batch effects (FDR<0.5; in case multiple batches were processed simultaneously). Samples were excluded when genotype coverage was <80%, or heterozygosity was >5SD above the mean (risk of contamination). Quality-controlled genotypes were phased using Eagle[4] and imputed with MaCH[5,6] using the trans-ethnic reference sample (v1.1) from the Haplotype Reference Consortium[7]. High quality variant sets from the quality control procedure were selected to compute individual's genetic principal components representing loadings along the 20 first eigenvectors of the pairwise genetic covariance matrix of a sub-sample of unrelated individuals from the HRC panel. Lastly, variants with INFO < 0.8 or MAF < 0.01 were excluded and samples with < 75% imputation confidence were set to missing, whereas the remaining samples were converted to best guess hard allelic dosages.

Blood samples of MoBa participants were obtained from both parents during pregnancy and from mothers and children (umbilical cord) at birth[8]. We obtained quality controlled and imputed genotyping data as described in the MoBaPsychGen pipeline v.1 by Corfield & Frei *et al*[9]. No additional quality control was performed.

**Loci and genes of original GWAS and condFDR summary statistics**

We defined genome-wide significant loci (see *Locus definition*) for the original GWAS as $p < 5\times10^{-8}$ and conditioned summary statistics as FDR < 0.05. We used the standard ways of reporting significant findings for considered analyses to keep consistency and facilitate comparison with previously reported results. However, we acknowledge that these criteria are not directly comparable, as discussed in the condFDR methods descriptions[10–12]. Since the probability of replicating these original and conditioned loci at genome-wide significance is low due to the limited statistical power of replication GWAS, we first tested for en masse sign concordance of effect direction in independent summary statistics, as is in line with previous literature[13,14]. All summary statistics that were used to look up lead SNPs and test for sign concordance are listed in Supplementary Data 12. An exact binomial test was used to test the null hypothesis that directions of effect were randomly distributed (*prob.*=0.5), given the total number of variants and the number of variants with concordant effects.

Using the genome-wide significant loci from the condFDR and original summary statistics, we used FUMA[15] (https://fuma. ctglab.nl/) to positionally map SNPs to genes. The resulting gene lists were used for hypergeometric test-based tissue enrichment analyses (as implemented in FUMA), to test the tissue-specificity of the input genes for differentially expressed (upregulated) genes in each of the 54 tissues available in GTEx (v8)[16]. Bonferroni correction was applied for the number of tests performed ($\alpha = 0.05 / (54 \text{ tissues} \times 2 \text{ gene lists} \times 5 \text{ disorders}) = 9.26\times10^{-5}$).

### Supplementary Results

*Comparison to previous studies*

For the current study, we used data from three previous studies that applied MOSTest on single neuroimaging modality phenotypes previously: functional MRI-derived phenotypes as used in Roelfs *et al*[17], structural MRI-derived phenotypes as used in van der Meer & Frei *et al*[18], and diffusion MRI-derived phenotypes Fan *et al*[19]. The number of loci we identified for each modality (640 sMRI, 44 fMRI, 562 dMRI, 851 multimodal at $p < 5\times10^{-8}$) cannot be compared one-to-one with these publications, because of some difference in analytical choices between the current and the previous studies. First, the previous publications considered phenotypes from each modality to consist of subgroups (structural phenotypes can be divided into measures of cortical thickness, cortical surface area, and subcortical volume; diffusion phenotypes can be divided into N0, ND, and NF principal components; functional phenotypes can be divided into within-network temporal variance, and between-network correlations) and analyzed these subgroups of phenotypes into separate MOSTest analyses. In contrast, we were interested in the genetic signal specific and shared across modalities, so we included all phenotypes derived from the same MRI modality into one MOSTest analysis. Second, the sample sizes are generally larger in the current study, because new data have been released since publication or the previous study used a within sample replication (UKB) instead of our independent sample replication procedure (ABCD). Third, the analysis pipeline used in the current study needed slight harmonization of the pipelines used in the previous studies, such as dimensionality reduction (diffusion, Supplementary Fig. 10), standardizing the covariates used or filtering on significantly heritable phenotypes prior to multivariate GWAS (see Methods).

*Summary statistics used for conditional FDR*

In the original schizophrenia article, the authors report 287 genome-wide significant loci based on the primary trans-ancestry GWAS ($N_{case}$=74,000; $N_{control}$=101,000). Since our locus definition procedure is slightly different, and uses a European reference panel (1000Genomes), we report 257 loci in these summary statistics. Additionally following requirements of the condFDR method, we had to limit our analyses to the European subsample ($N_{case}$=53,000; $N_{control}$ =77,000; $n_{loci}$ =177) and exclude the TOP sample to prevent sample overlap ($N_{case}$=51,000; $N_{control}$=68,000; $n_{loci}$=162).

We also used summary statistics for major depression disorder published by Levey et al., though with exclusion of several samples. The original summary statistics are a meta-analysis of the Million Veteran Program, PGC (Wray et al.), 23andMe, UKB and FinnGen. However, the 23andMe summary statistics are not publicly available, we already use FinnGen summary statistics for our condFDR replication phase and could not allow sample overlap with our UKB-based MOSTest summary statistics. This decreased the sample size by 66% and therefore also the number of loci from the reported in the original publication ($n_{loci}$= 178) to $n_{loci}$=20.

In the publicly available autism spectrum disorder summary statistics 2 genome-wide significant loci are reported (chr20 and chr8; $N_{case}$=18,000; $N_{control}$ =27,000). In the accompanying article, the additional loci reported are discovered in a combined analysis with a follow-up sample ($N_{case}$ =2,000; $N_{control}$ =142,000) that is not publicly available.

The loss of 5 loci (64 in the original article at $N_{eff}$=101,962, we report 49 at $N_{eff}$=93,662) for bipolar disorder is due to excluding the UKB and TOP sample to prevent sample overlap. There was no sample overlap with the samples used for the original attention deficit hyperactivity disorder GWAS, and we report the same number of loci as the original article ($n_{loci}$=12).

*Conditional FDR and original GWAS loci and genes*

The genome-wide significant loci identified in the original GWAS and in the condFDR summary statistics are displayed in Supplementary Data 13. Note that the number of significant loci from the original GWAS may be lower than the number presented in the original articles, since we relied on publicly available data (e.g. excluding 23andMe) and exclude some additional cohorts to prevent sample overlap (with e.g. UKB, see Supplementary Results 2). The sign concordance of the original lead SNPs and condFDR lead SNPs using independent disorder GWAS summary statistics[20–22] can be found in Supplementary Data 14. We positionally mapped condFDR loci to genes (528 genes for MDD, 2,416 for SCZ, 116 for ASD, 1,162 for BD, 122 for ADHD; Supplementary Data 15) and subsequently used these genes as input for hypergeometric test-based tissue enrichment analyses, both as implemented in FUMA. The same test was applied on genes mapped from the original GWAS loci (60 genes for MDD, 498 for SCZ, 5 for ASD, 186 for BD, 14 for ADHD; Supplementary Data 15). Supplemental Fig. 9 shows the results of the enrichment test for gene-sets with upregulated gene expression in brain and bodily tissues (Supplementary Data 16).

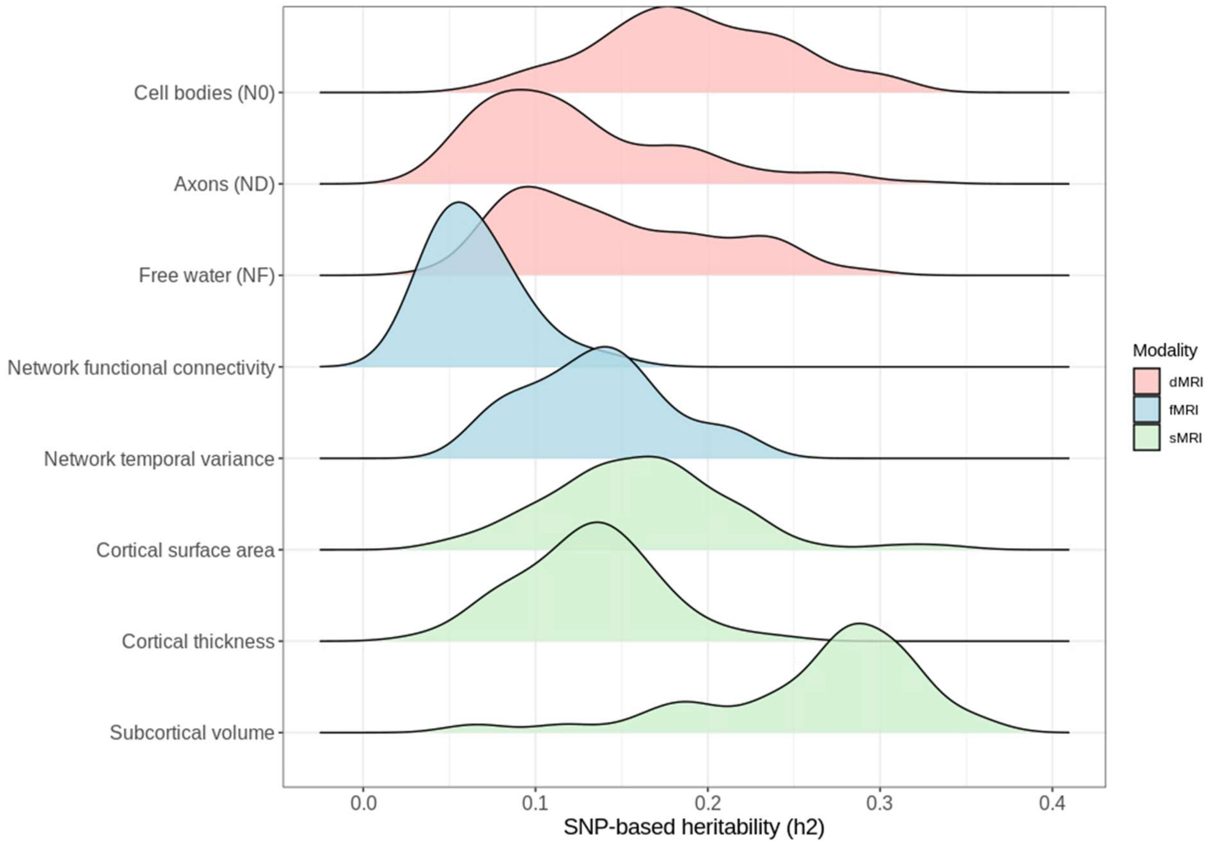

*Supplementary Figure 1*. Heritability of all phenotypes coming from three different modalities (dMRI = diffusion MRI, fMRI = functional MRI, sMRI = structural) after filtering on nominal significance. Estimates are obtained using LDSC Regression (Methods).

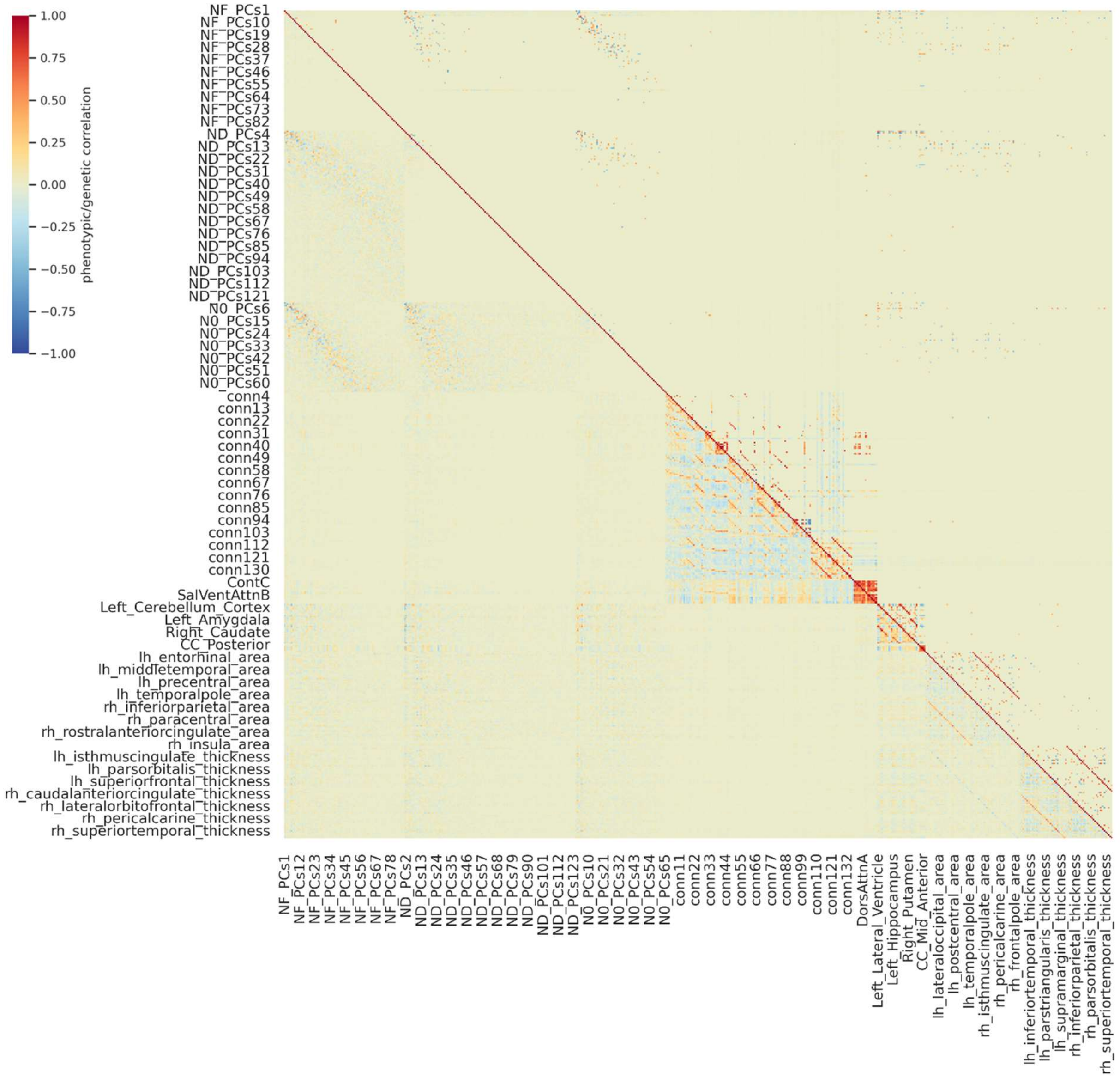

*Supplementary Figure 2.* Phenotypic (lower triangle) and genetic (upper triangle) correlation matrix between all phenotypes used in this study. Insignificant correlations (Bonferroni corrected $p > 0.05$) are set to zero for visualization purposes. Note that correlations within each dMRI-derived tissue metric are zero, because these phenotypes represent the orthogonal principal components within every metric.

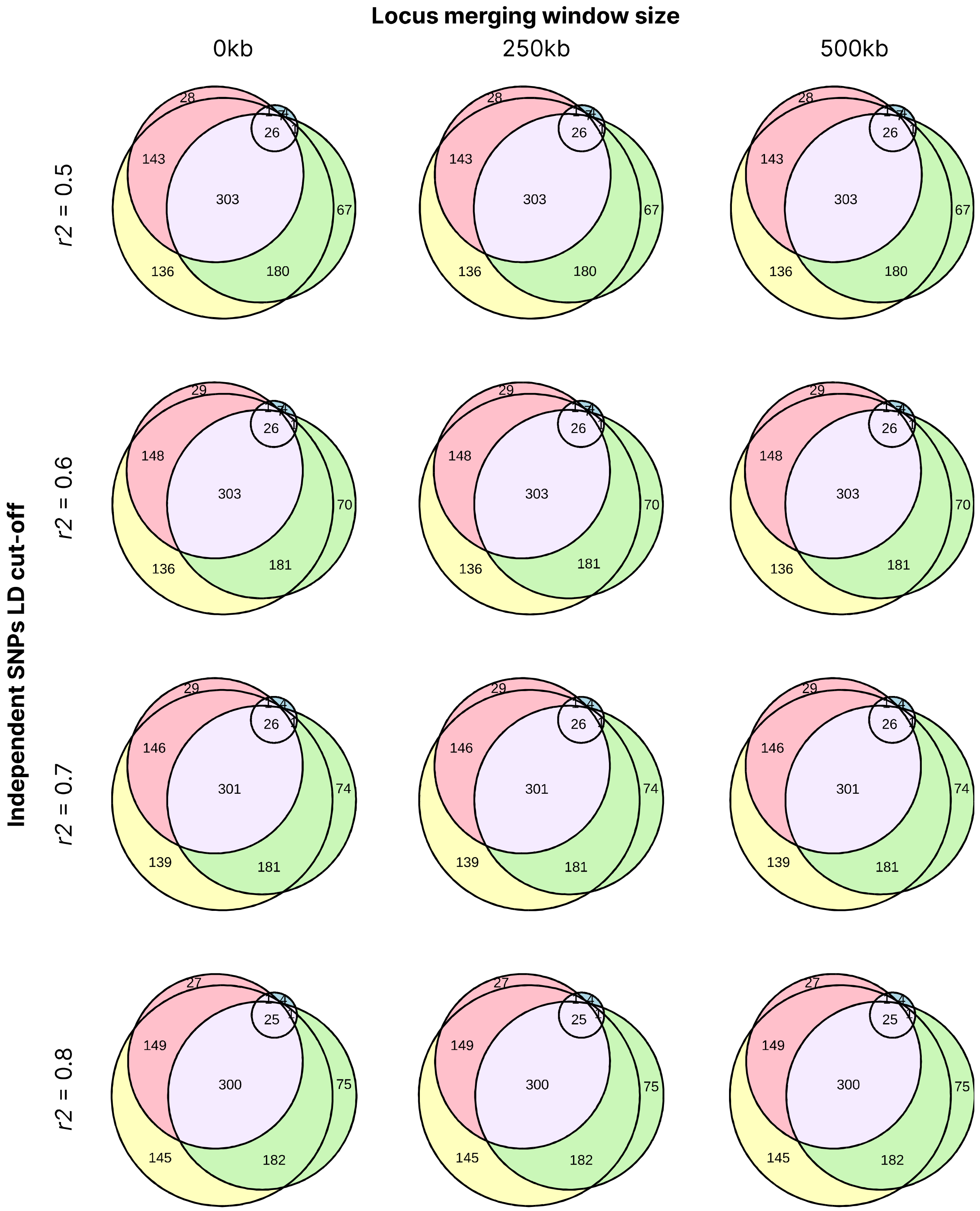

*Supplementary Figure 3.* Sensitivity analyses for the effect of locus definition parameters (window size and $r^2$) on locus overlap. Varying the settings has minute effects on the number of (overlapping) loci.

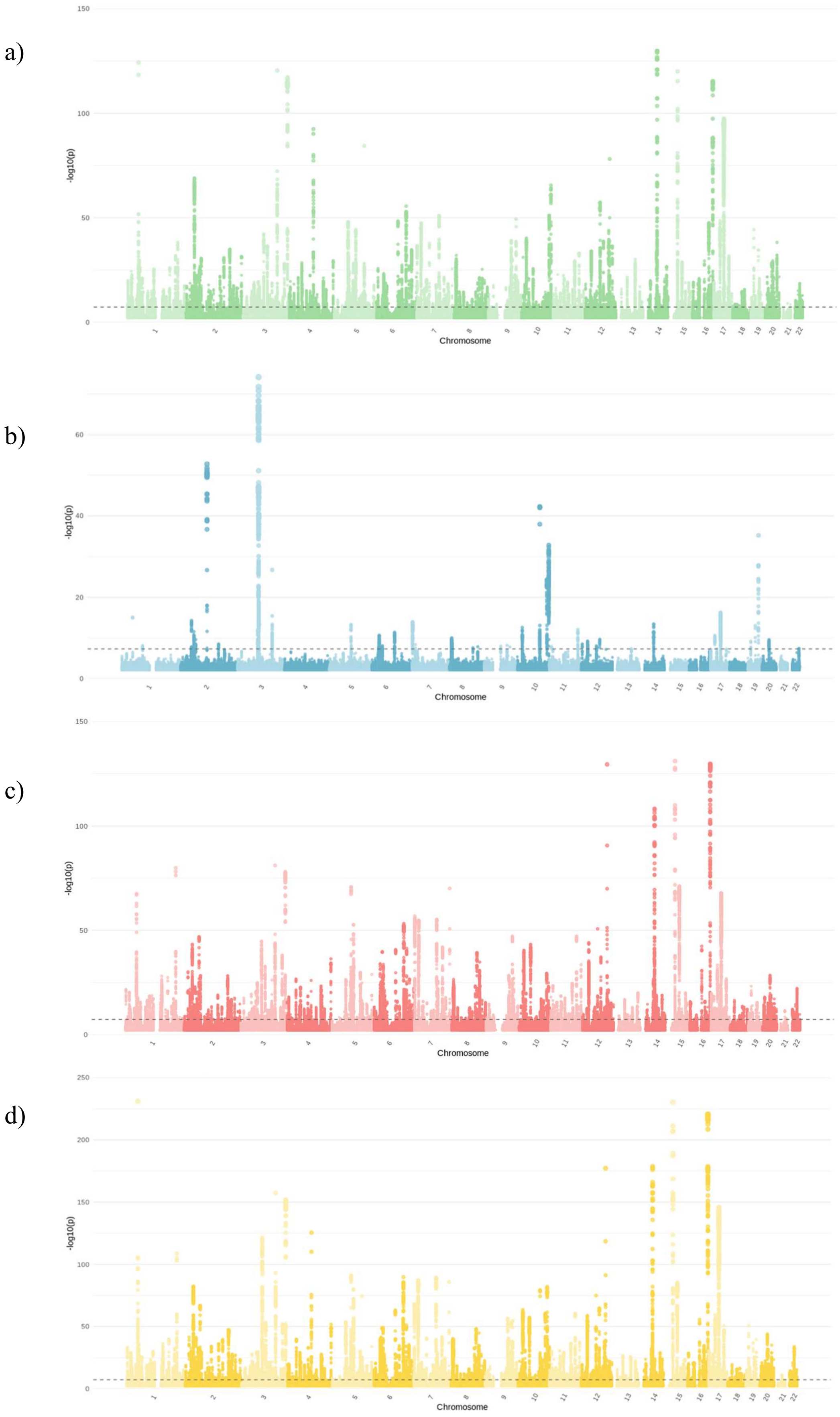

*Supplementary Figure 4.* Manhattan plots of the multivariate GWAS results. We ran MOSTest for all phenotypes coming from a) structural MRI, b) functional MRI, c) diffusion MRI, or d) all combined.

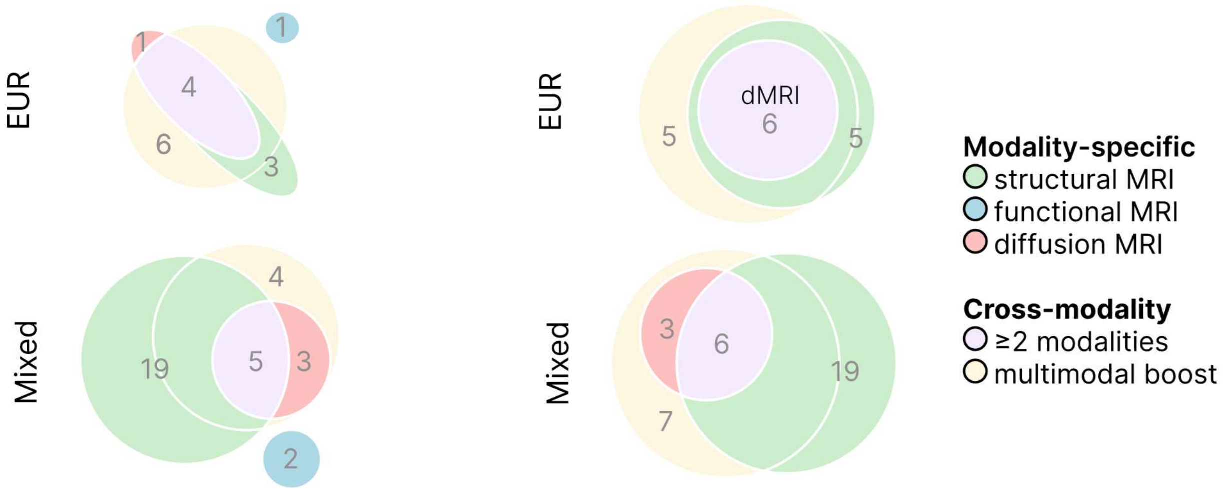

*Supplementary Figure 5*. Overlap of genome-wide significant loci (left) and genes (right) observed across neuroimaging modalities in single-modality and joint multimodal replication analyses in the ABCD sample.

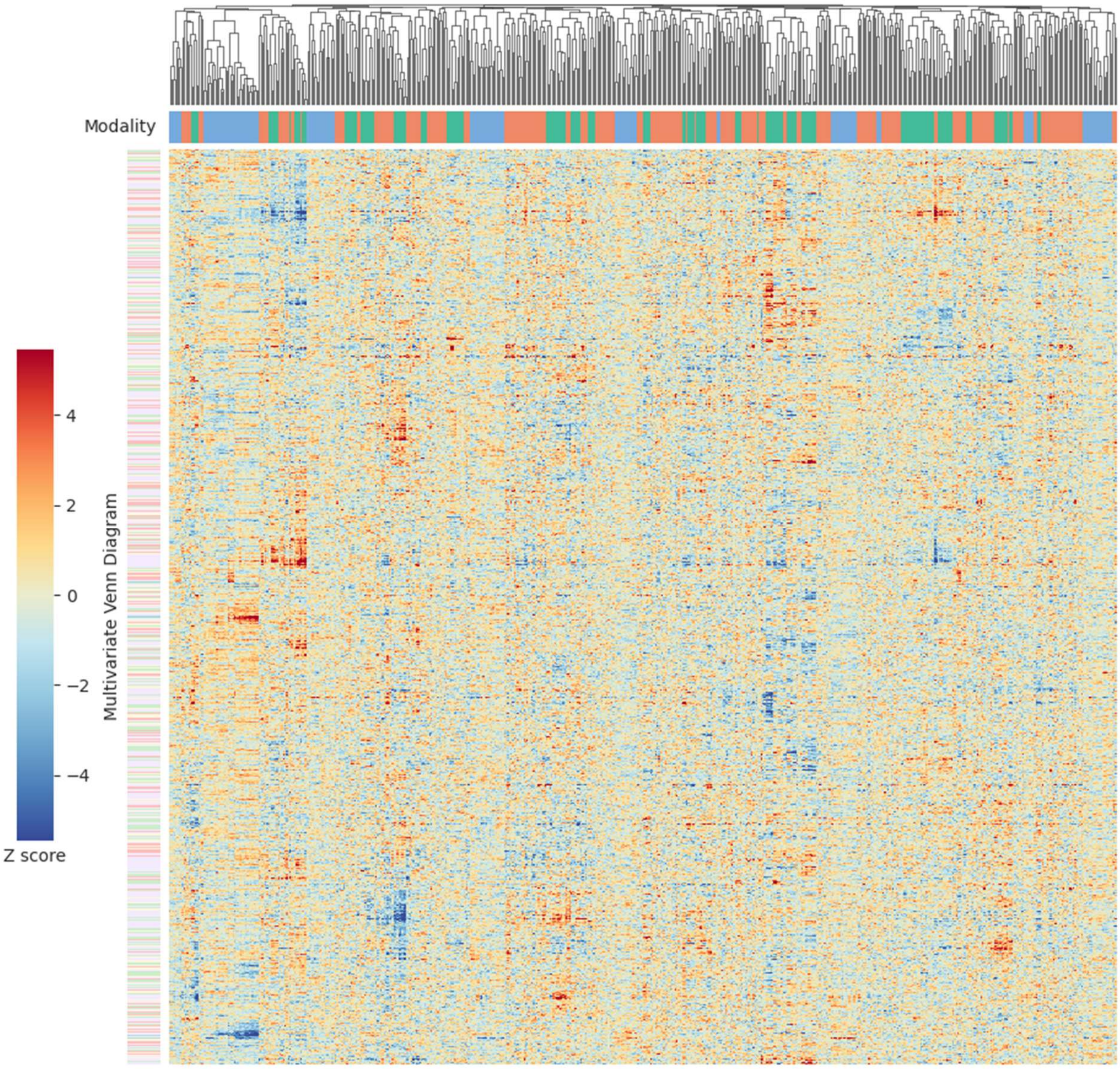

*Supplementary Figure 6.* Univariate *p*-values underlying all lead SNPs identified through multivariate GWAS in MOSTest. Lead SNPs are categorized by their location in the Venn diagram (Figure 1a in main text) on the y-axis, phenotypes are clustered on the x-axis. Hierarchical clustering was applied (Methods).

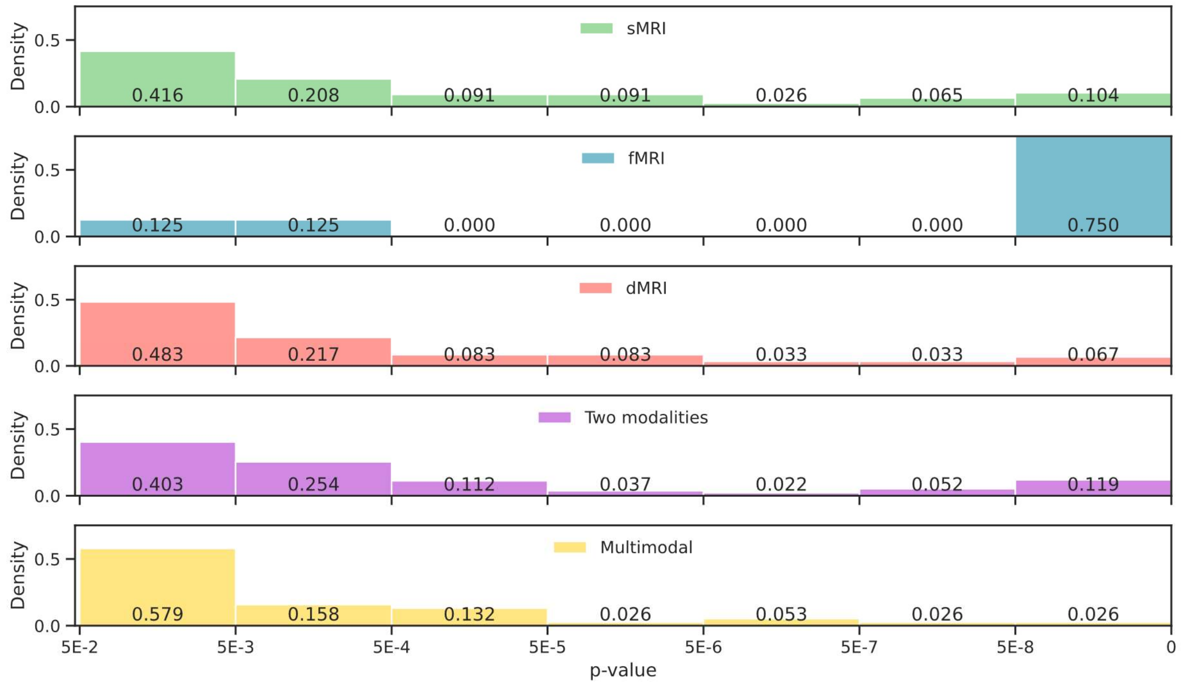

*Supplementary Figure 7.* Minimal univariate *p*-value underlying all lead SNPs identified through multivariate GWAS in MOSTest. Lead SNPs are categorized by their location in the Venn diagram (Figure 1 in main text). Please note the relative low number of fMRI-modality lead SNPs.

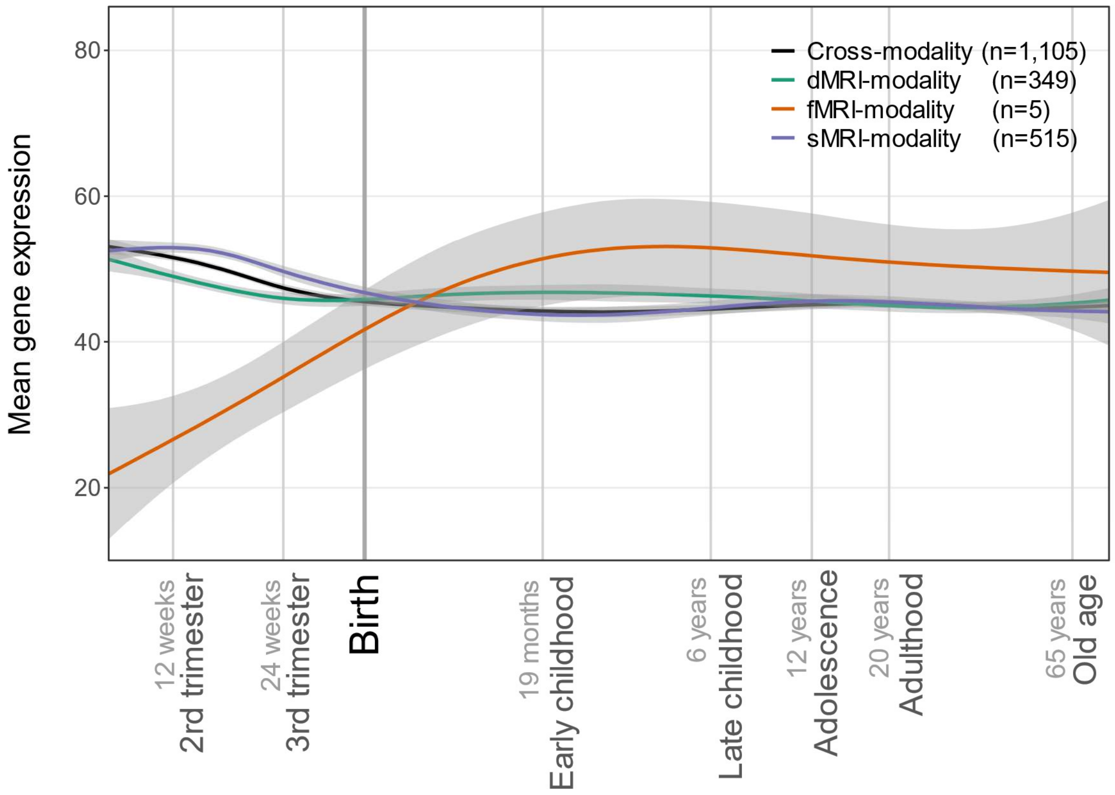

*Supplementary Figure 8.* Mean-normalized expression (y-axis) of cross-modality and single-modality genes over developmental timepoints (x-axis; log10 scale). Gray shading indicates 95% confidence intervals. Note that for fMRI-modality genes *n*=5, so the observed pattern should be interpreted with caution.

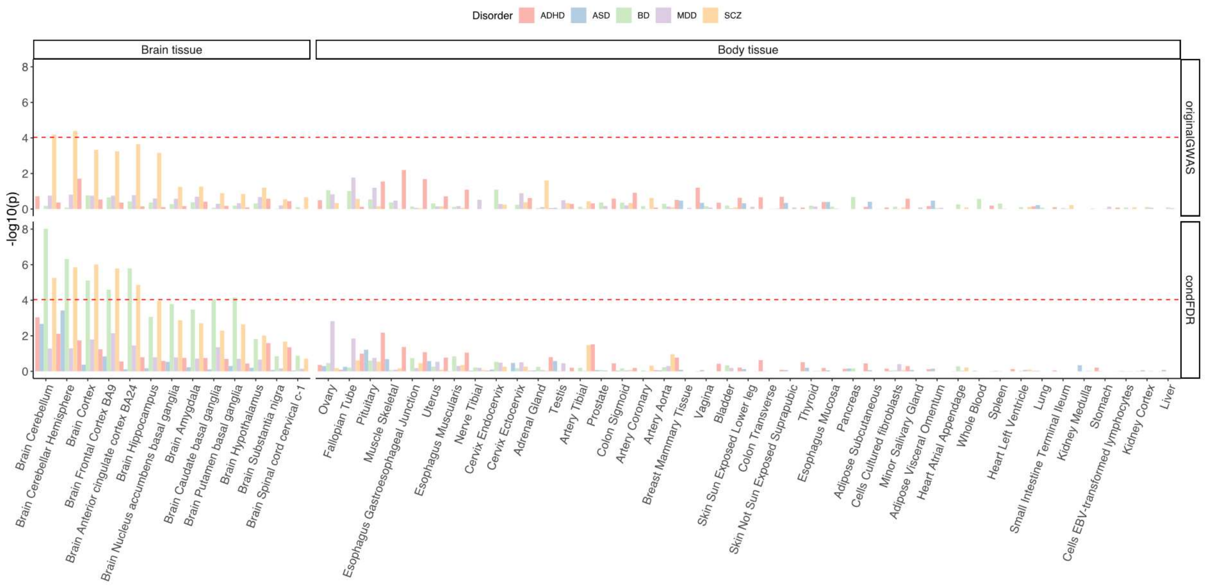

*Supplementary Figure 9.* Enrichment of disorder genes (positionally mapped from the original disorder GWAS loci *vs* conditional FDR loci) in tissue-specific gene-sets with upregulated expression (Supplementary Data 16). The red line indicates Bonferroni correction for the number of tests ($\alpha = 0.05/540 = 9.26\times10^{-5}$).

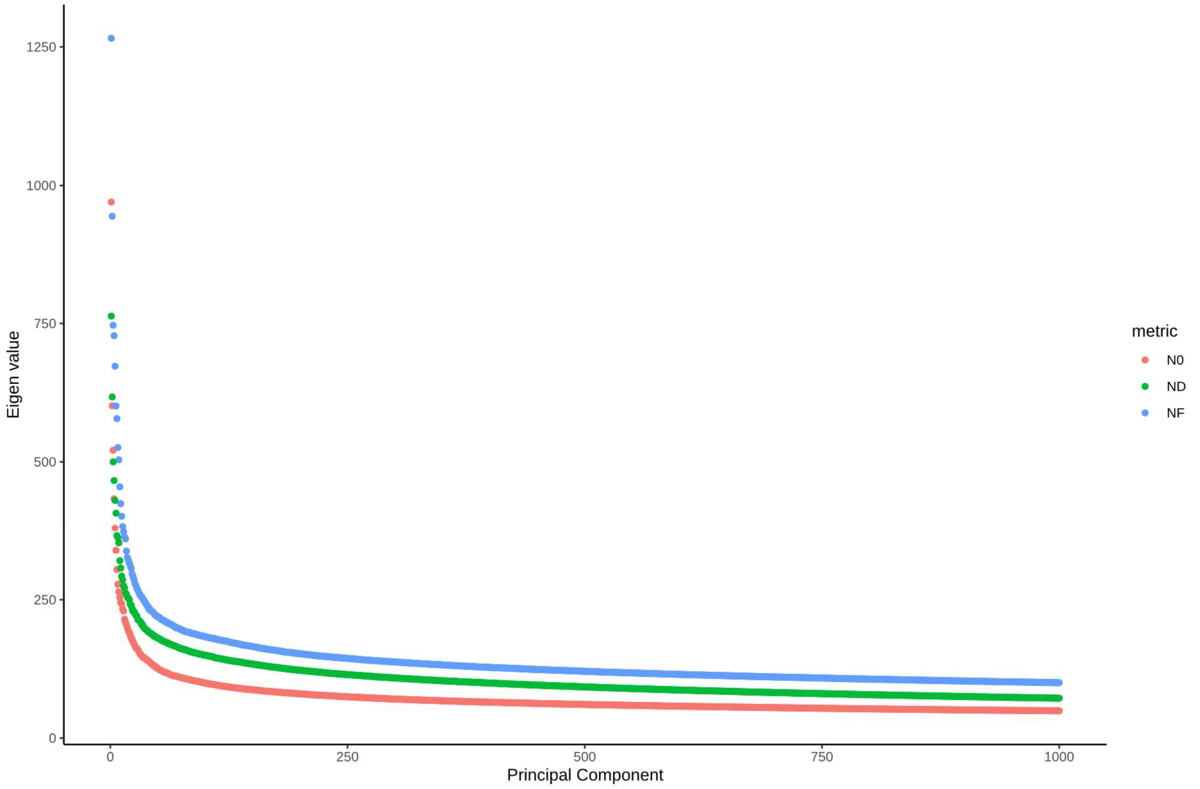

*Supplementary Figure 10.* Scree plot for the diffusion MRI-derived phenotypes as used by Chun *et al* (2022). For reasons of dimensionality reduction, we estimated the “elbow” of the eigenvalues for each dMRI-derived measure (ND, NF, N0) and used those PCs in our study.
